# Targeted Time-Varying Functional Connectivity

**DOI:** 10.1101/2024.08.07.606804

**Authors:** Sonsoles Alonso, Luca Cocchi, Luke J. Hearne, James M. Shine, Diego Vidaurre

## Abstract

To elucidate the neurobiological basis of cognition, which is dynamic and evolving, various methods have emerged to characterise time-varying functional connectivity (FC) and track the temporal evolution of functional networks. However, given a selection of regions, many of these methods are based on modelling all possible pairwise connections, diluting a potential focus of interest on individual connections. This is the case with the hidden Markov model (HMM), which relies on region-by-region covariance matrices across all pairs of selected regions, assuming that fluctuations in FC occur across all investigated connections; that is, that all connections are locked to the same temporal pattern. To address this limitation, we introduce *Targeted Time-Varying FC* (T-TVFC), a variant of the HMM that explicitly models the temporal fluctuations between two sets of regions in a targeted fashion, rather than across the entire connectivity matrix. In this study, we apply T-TVFC to both simulated and real-world data. Specifically, we investigate thalamocortical connectivity, hypothesizing distinct temporal signatures compared to corticocortical networks. Given the thalamus’s role as a critical hub, thalamocortical connections might contain unique information about cognitive processing that could be overlooked in a coarser representation. We tested these hypotheses on high-field functional magnetic resonance data from 60 participants engaged in a reasoning task with varying complexity levels. Our findings demonstrate that the time-varying interactions captured by T-TVFC contain task-related information not detected by more traditional decompositions.

## 1. INTRODUCTION

Functional connectivity (FC) has become a key metric for studying brain communication, capturing statistical dependencies between brain signals over time (Biswal et al., 1995; Smith et al., 2013), and providing insights into the coordinated activity of brain regions (Smith et al., 2009). Time-varying FC extends this framework by capturing the dynamic nature of these interactions, recognizing that brain networks might not be static, but rather exhibiting fluctuations in connectivity patterns over time (Breakspear, 2017; Calhoun et al., 2014; Zalesky et al., 2014). This understanding of dynamic changes is important for characterising how brain networks support various cognitive functions, including memory, attention, and decision-making (Cocchi et al., 2013; Kucyi et al., 2017; Menon and D’Esposito, 2022; Wang et al., 2021). To better capture such fluctuations, many techniques have been developed and refined (Iraji et al., 2021; Lurie et al., 2020; Xie et al., 2019).

A common approach for modelling time-varying FC in neuroimaging data is the use of state-based models, such as the hidden Markov model (HMM). These models characterize brain timeseries as temporal sequences of states, where each state is associated with a distinct pattern of FC, thus facilitating the identification and analysis of dynamic changes in brain connectivity over time (Vidaurre, 2021; Vidaurre et al., 2021; Vidaurre et al., 2017). To do so, current methods such as the HMM —but also clustering algorithms based on sliding window estimates (Allen et al., 2014) or approaches built on the graphical lasso (Cai et al., 2019; Monti et al., 2014)— rely on region-by-region covariance matrices that include all possible pairwise interactions. This inherently limits the ability to focus on specific connections of interest and assumes that all aspects of brain connectivity vary in a temporally consistent manner, i.e. all at once. We suggest that this simplification may overlook the temporal subtleties of specific connections, which may be important to understand specific aspects of cognition. Furthermore, methods that can theoretically handle non-square matrices, such as those based on windowed-correlation matrices (Hussain et al., 2023), still face the challenge of arbitrary window length selection, which becomes particularly problematic when attempting to capture rapid changes within short trial periods. To address these limitations, we introduce a novel, windowless approach called *Targeted Time-Varying FC* (T-TVFC), which extends the conventional HMM framework to target specific sections of the connectivity matrix. Using simulations, we demonstrate the advantage of focusing on specific connections when these are the ones that matter to our scientific question.

One example of a relevant subset of connections is the thalamocortical system (Shine et al., 2023). Here, we hypothesise that the connections between the thalamus and cerebral cortex may exhibit temporal signatures that are separate from, for example, other corticocortical processes. For instance, as reasoning demands increase, specific thalamocortical pathways are likely to support the reconfiguration of interactions between spatially distributed networks, which may be important for higher-order cognition (Cocchi et al., 2014; Hearne et al., 2017; Parkin et al., 2015). Isolating the dynamics of these thalamocortical connections may help us dissect the nature of these processes. In contrast, coarser network analyses of dynamics based on full connectivity matrices across all selected regions might shroud these dynamics in favour of other processes that are not central to our question of investigation. To explore how the thalamus interacts with the cortex dynamically at a large scale, and showcase the utility of T-TVFC, we examine a task with varying cognitive demands. Specifically, we used high-density (7T) functional magnetic resonance imaging (fMRI) data from 60 participants engaged in a non-verbal relational reasoning task with varying levels of complexity (Birney et al., 2006; Hearne et al., 2017). We show that the proposed model effectively captures fluctuations in thalamocortical connectivity, revealing a stronger relationship to task difficulty than models of connectivity across all regions.

## 2. MATERIALS AND METHODS

### 2.1. MODELS OF CONNECTIVITY DYNAMICS

T-TVFC is a time-varying model specifically designed to capture dynamic interactions between two sets of brain regions. Built upon the HMM framework, T-TVFC identifies recurrent states characterised by specific connectivity patterns between these two sets. Unlike the conventional HMM, where states represent connectivity matrices across all pairs of *p*×*p* regions, T-TVFC states represent connectivity matrices across specific pairs of *p*×*q* regions, where *p* is the number of regions in one set, and *q* in the other; these sets can be overlapping or non-overlapping. This targeted approach allows for a more focused analysis of hypothesis-driven neural interactions. The general HMM framework, the conventional HMM and the proposed T-TVFC model are described next and illustrated in **Figure 1**.

**Figure 1.**
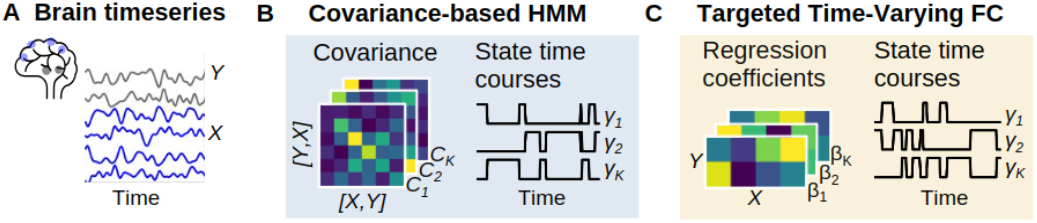
Schematic representation of the methodological approach. **A** Two sets of brain timeseries (*X* in blue and *Y* grey) representing the input data for the models. **B** The covariance-based HMM captures pairwise co-variation collectively across all regions in Z (i.e., [*X, Y])*, by discretizing them into a set of *K* states; each state is associated with a probability of activation per time point, represented as HMM state time courses. **C** The T-TVFC captures changes in the interaction between timeseries *X* and *Y*; specifically, each state corresponds to a distinct set of β coefficients linking *X* to *Y*, without modelling the connections within *X* or within *Y*.

#### 2.1.2. The HMM framework

The HMM is a probabilistic model employed for the analysis of timeseries data, and other types of sequential data such as genetics. When applying to brain data, the HMM represents neural activity as a sequence of *K* latent (or hidden) states, each representing a unique pattern of activity and FC. Each state is characterized as a probability distribution. At each time point, the inference of the model outputs *K* probabilities of activation, corresponding to the likelihood of the data being generated by each state. These probabilities are also influenced by the probabilities of the previous time point. Mathematically, the state dynamics can be described as follows:

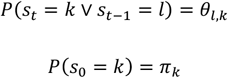

Where θ_*l*,*k*_ is the probability of transitioning from state *l* to state *k;* and π_*k*_ is the probability that each segment of the timeseries (e.g. session) starts with state *k*. Given this, we define the estimated state probabilities (referred to as state time courses) as γ_*t*,*k*_ ≔ *P*(*s*_*t*_= *k* ∨ *s*_<*t*_,*s*_>*t*_,Z.), where *Z* is the data and *s*_<*t*_,*s*_>*t*_ represent, respectively, the state time course before and after time point *t*.

This is a general description of the HMM framework, which can be instantiated into various specific models depending on which probability distribution we choose to describe the states.

##### 2.1.1.1. Conventional (covariance-based) HMM

In the most common model for continuous data, the states are represented as Gaussian distributions where the covariance matrix of each state, Σ^*k*^, captures the covariation across all pairs of regions. To focus on changes in FC, the mean parameter of the Gaussian distribution, which encodes the average activity per region for state *k*, can be set to zero. Mathematically, if state *k* is active at time point *t*, the data is distributed

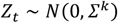

This representation is equivalent to a Wishart-distribution-based HMM (Vidaurre et al., 2021).

##### 2.1.1.2. Targeted Time-Varying FC Model

As shown in Vidaurre et al. (2025), we can generalise the Gaussian model to the following:

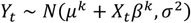

Where *Y*_*t*_ and *X*_*t*_ represent two sets of timeseries (the response and the regressor of a regression model); β^*k*^ is a matrix of regression coefficients linking *X* to *Y* when state *k* is active; μ^*k*^ is the mean parameter of the distribution; and σ^2^ represents a diagonalised vector of variances that is not state-specific. If we, for example, assume that *Z*_*t*_ contains both *Y*_*t*_ and *X*_*t*_ concatenated, and fix μ^*k*^=0, this model expresses an off-diagonal submatrix of the full covariance represented in the covariance-based HMM, emphasising the specific connections of interest.

##### 2.1.1.3. Reproducible models

To enhance the reproducibility of model results from both the conventional HMM and T-TVFC, we implemented a strategy to reduce the variability introduced by random initializations. This variability is commonly observed any model which estimation of parameters is performed through mathematical optimisation; this is the case of the HMM, where the optimization starts with random initial values, such that the solution may depend on this initialisation. To address this, we ran the model five times and selected the best-ranked HMM using a quantitative metric known as free energy, which combines fitness and model complexity (Alonso and Vidaurre, 2023). To ensure reproducibility, we repeated this process 100 times and assessed result similarity across repetitions. We quantified similarity using the Hungarian algorithm (Munkres, 1957) for state alignment, calculating similarity based on a cost matrix reflecting state dissimilarity. The similarity metric similarity (i, j) = 1 – (cost / max cost) indicates the degree of consistency between runs, ranging from 0 to 1. Across all models in this study, we observed high similarity between repetitions, with average similarities of 0.99 for the T-TVFC and 0.96 for the conventional HMM (see histograms in **Figure S1**). This indicates that selecting the best-ranked run (i.e., with the lowest free energy) from five model runs provided reproducible results.

#### 2.1.2. Sliding windows

We evaluated T-TVFC against a sliding-window approach for capturing dynamic FC. In this method, Pearson’s correlations between pairs of regions of interest were calculated within a fixed-length window, which slides one time point at a time. K-means clustering was then applied to the resulting sequence of sliding-window correlation (SWC) matrices, identifying clusters that allegedly represent distinct FC states that emerge throughout the scanning session, along with the timing of state transitions (Allen et al., 2014). A critical challenge here is selecting an optimal window length that adequately balances sensitivity to transient connectivity fluctuations with stability for capturing broader connectivity patterns (Hutchison et al., 2013). Recommended window lengths widely range from 20 to 100 seconds (Preti et al., 2017; Sakoğlu et al., 2010; Vergara et al., 2019). To keep the comparison agnostic, we implemented the sliding-window approach over a range of window sizes to characterize FC states for all *p*×*p* region pairs s well as the targeted *p*×*q* pairs.

### 2.2. SIMULATED DATA

We first used synthetic data to demonstrate the efficiency of the T-TVFC approach compared to the conventional (covariance-based) HMM in capturing the temporal fluctuations of specific connections. The simulations were designed such that the task-relevant information was contained in specific connections only. In this case, as we anticipated, the T-TVFC would more closely captures the task-relevant information. In contrast, the covariance-based HMM, which considers the joint statistical properties across all regions indistinctly (i.e. including variances and all cross-covariances) is less sensitive.

#### 2.2.1. Data Description

We synthetically generated two sets of timeseries, *X* and *Y*, which relate to each other through a regression model where *X* correspond to the regressors and *Y* to the responses. The nature of such regression had a changing relationship over time. With respect to the regressors, *X* is a matrix with *p* channels sampled under two different scenarios. In the first scenario, *X* is sampled from a covariance-based HMM, such that the covariance across regions changes over time according to state time courses that were also sampled randomly. In our tests, the number of regions in *X* ranged from 1 to 25; we sampled 1,000 trials (500 trials per condition) of 100 time points each trial. In the second scenario, *X* comprised timeseries from 11 cortical networks from resting-state 7T-fMRI data, obtained from a larger dataset, used in the real data experiments (see below). Here, the number of predictor variables, *p*, was set to vary from 1 to 11, as we randomly selected a subset of these cortical network timeseries.

In either case, we then sampled *Y*, with one (*q*=1) channel only, using another HMM model (of note, with different state time courses than those of the generation of *X* in the first scenario just described). This HMM model was a T-TVFC model, with condition-specific and state-dependent regressors. The state time courses of this HMM were generated for *K*=5 states, such that states followed one another in sequence for all trials (i.e. from state 1 to *K*), and stayed active for approximately equal time; however, we introduced some temporal variability across trials regarding the onset and offset of the states. Each state was endowed with a randomly generated matrix of (*p* by *q*) regression coefficients β, defining distinct *X*-to-*Y* relationships. To introduce variability between task conditions, we manipulated the relationship between *X* and *Y* for a specific state. In particular, we induced differences in β for state 2 by adding independent normally-distributed noise for each condition. Finally, we added normally distributed noise to *Y*, to achieve various degrees of signal-to-noise ratio; specifically, the standard deviation of the noise was a factor ranging from 0.001 to 0.2 of the standard deviation of the signal (which is given by *X*β).

### 2.3. REAL DATA

We used 7T fMRI data with a TR of 0.576 seconds, collected from 60 participants during three 10-minute task sessions (Hearne et al., 2017). The participants were tasked with a non-verbal relational reasoning paradigm comprising problems of varying difficulty levels categorized as null, binary, ternary, and quaternary based on relational complexity. Each condition consisted of 36 trials totalling 2,160 trials across subjects. As outlined in Hearne et al. (2017), task performance decreased with increasing complexity, with increased errors and response time attributed to task demands rather than temporary lapses in attention or disengagement. Correct responses represent 96% for binary trials, 87% for ternary trials, and 65% for quaternary trials. Participants demonstrated a success rate of 96% on null trials, further indicating sustained attentiveness and task engagement.

#### 2.3.1. Data Description

Acquisition and processing details can be found in Shine et al. (2019). In brief, echo planar images were obtained with a multiband sequence. High-resolution anatomical images were captured for preprocessing. Data preprocessing included realignment, reorientation, and co-registration to T1 images. Segmentation and DARTEL were used for signal estimation and spatial normalization. The aCompCor method was used to remove residual non-neural signals. Participants with excessive head movement (> 3 mm in over 5% of the volumes) were excluded. No temporal filtering was applied. After pre-processing, the mean regional timeseries for each ROI was computed by averaging the voxel time series within each ROI. These timeseries were extracted from 333 pre-defined cortical parcels using the Gordon atlas (Gordon et al., 2016), and from 31 thalamic regions using the Morel atlas (Niemann et al., 2000). Only 17 of the 31 regions were included in further analysis as the limited spatial resolution of BOLD data (approximately 2 mm^3^ isotropic voxels) prevented us from modelling the lateral and medial geniculate nuclei and the intralaminar nuclei with sufficient detail. Following the extraction of the timeseries, Finite Impulse Response (FIR) modelling was utilized to estimate the haemodynamic response function (HRF). Unlike the conventionally used canonical HRF, FIR models capture the haemodynamic response at each time point without assuming a predetermined shape. To execute this, a distinct design matrix was created for each participant/session, representing each trial (categorized by Complexity [e.g., binary, ternary, or quaternary] and Performance [e.g., Correct or Error]) as an FIR (i.e., with unique regressors for each TR within a trial).

Finally, to reduce data dimensionality, we applied principal component analysis (PCA) to identify the major activation patterns within each of the 11 functional networks, as delineated in the adopted Gordon et al. (2016) atlas: Auditory (AUD), Cingulopercular (CO), Cinguloparietal (CP), Dorsal Attention (DAT), Default Mode (DM), Frontoparietal (FP), Retrosplenial Temporal (RT), Sensorimotor (SM), Salience (SAL), Ventral Attention (VAT) and Visual (VIS). Cortical regions not associated with any of these networks were excluded. For the 17 thalamic timeseries, 12 were grouped into the anterolateral (AL) nuclei, three into the pulvinar (Pu) nuclei and one into the medial dorsal (MD) nucleus. The first principal component (PC) was extracted for each group, except for the MD, which consists of only one region. The first principal component (PC) of each cortical network and thalamic nucleus captured a substantial portion of the variance, typically exceeding 70%. While the explained variance showed a slight increase with task complexity (**Figure S2**), statistically significant differences across task conditions were only observed for the DM and DAT networks. **Figure S3** presents the averaged PC time series of each cortical network and thalamic nucleus across subjects, stratified by task complexity. The signal-to-noise ratio (SNR) for most networks was relatively low, with both positive and negative values observed across task conditions (**Figure S4**). This may be due to trial-to-trial variability and edge effects at the beginning and end of each trial.

### 2.4. EMPIRICAL ASSESSMENT

To assess the models’ ability to capture task-relevant information, we calculated classification accuracy in distinguishing task conditions over time using state time courses derived from each model. For each subject, the state time courses were epoched by trial onset and averaged across trials of the same condition, resulting in condition-specific mean trial-locked state time courses. At each timepoint *t*, these subject-level averages, arranged in a matrix with subjects as rows and conditions as columns, were used as predictors for classification. We used logistic regression for the simulated data (two conditions); and ordinal logistic regression for the real data, given the ordered nature of the conditions (binary, ternary and quaternary). All runs of the models were unsupervised, with no prior knowledge of task timings. We applied 5-fold subject-wise cross-validation, ensuring that subjects were not split between folds (i.e., a model tested on a given subject had not seen any trial of that subject during training). Within each fold, the predictive model was trained on data from four folds (training data) and used to predict task conditions for the remaining fold (test data). Classification accuracy was computed as the percentage of correctly classified task conditions across subjects in the test fold. This process was repeated for each fold, yielding an overall cross-validated accuracy score. A perfect classification corresponds to an accuracy of 1, while 1/N indicates random guessing among N conditions.

## 3. RESULTS

As described in *Methods*, we used both simulated and real fMRI brain data to demonstrate how targeting a specific set of connections can offer a more sensitive description of the brain dynamics in relation to a task or cognitive process. To evaluate the models’ capacity to capture task-relevant information, we measured their classification accuracy in distinguishing between the simulated or real task conditions of the trials.

### 3.1. SIMULATED DATA

We first validated the performance of the T-TVFC model compared to the conventional HMM in recovering the temporal fluctuations of the connections of interest using simulated data where the ground truth is known. Specifically, we set the data to have two conditions, and used the capacity to predict which condition each trial belongs to from the model output as a measure of the models’ quality. As described in the *Methods* section, two different scenarios were considered regarding the nature of *X*: one where *X* is synthetic; and the other where *X* was directly extracted from real data. Either way, Y was generated synthetically such that its relationship to *X* varied over time within the trial and differed between the two conditions within a specific time window. For each scenario, the analysis was conducted across a range of predictors in *X* (*p*) and levels of noise (σ). Each simulation configuration was repeated 100 times. For each of these, the T-TVFC model and the covariance-based HMM were run and evaluated. For the T-TVFC model, *X* and *Y* were treated individually; whereas for the covariance-based HMM, *X* and *Y* were concatenated channel-wise into a single matrix *Z*. Both models were endowed with *K*=7 states, i.e. they were over specified since the ground-truth model has 5 states; we made that choice to give the covariance-based HMM an additional chance to adapt to its own misspecification (since the ground-truth is a T-TVFC model). The inferred state time courses of each model were then used to predict trial condition over time using logistic regression. Since we had two conditions, the baseline accuracy is 0.5.

In the first scenario, where *X* is synthetic, the T-TVFC model outperformed the covariance-based HMM as expected. **Figure 2A** illustrates the accuracy scores over time for each model for one specific setting (σ = 0.1 and *p* = 5). As observed, T-TVFC effectively captures the task-induced changes in the *X*-to-*Y* interactions during the time points when the effect of the condition was present. In the other panels, the accuracy of each model is represented by the average accuracy over this time window. **Figure 2B** displays the mean accuracy of each model across 100 repetitions for various combinations of *p* (from 1 to 25) and σ (from 0.001 to 0.18) values. Here, the T-TVFC model consistently outperformed the covariance-based HMM across different values of *p* and levels of noise; see **Figure 2C** for an explicit depiction of the difference. Lower levels of noise and higher number of variables in *X* led to an increased advantage of T-TVFC. When *X* consists of only one variable and the levels of noise are minimal, both models perform similarly because global changes in connectivity closely resemble distinct, localised changes. Conversely, when *X* had more than one channel, the covariance matrices contain more task-irrelevant information, which challenges the covariance-based HMM’s ability to isolate and capture the specific *X*-to-*Y* relationship.

**Figure 2.**
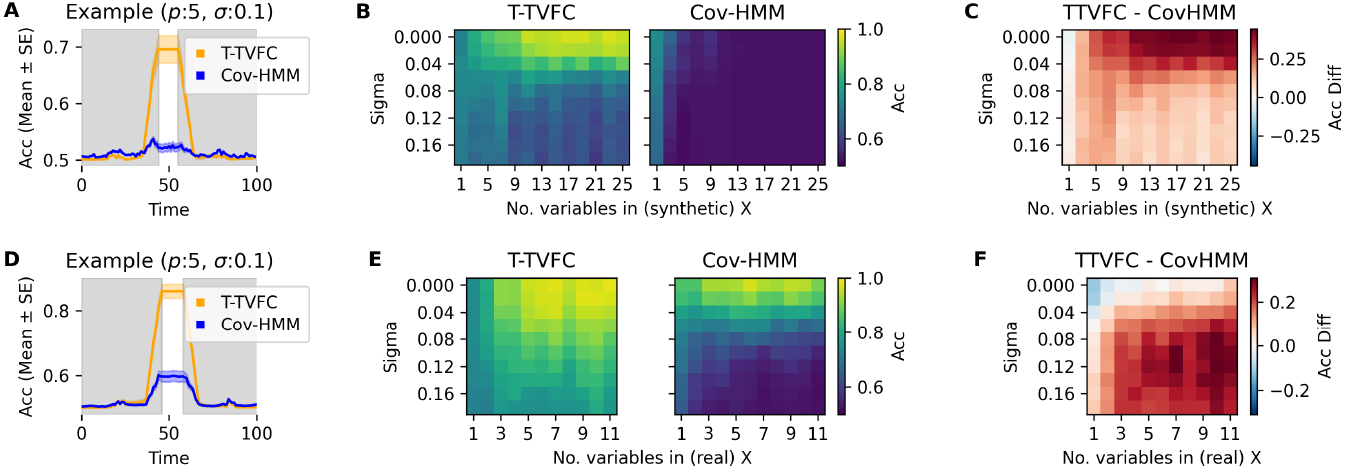
Model performance from simulations with synthetic (top) and real (bottom) predictors. **A, D** Example illustrating the performance of the T-TVFC (orange) compared to the covariance-based HMM (blue) over time for a specific parameter configuration (*p* = 5 and σ = 0.1). The shaded areas represent the standard error around the mean accuracy across 100 repetitions. The white patch indicates the period where the simulated task-dependent state (state 2) was active. **B, E** Mean accuracies for each model computed during the interval when there are differences between conditions, across *p* and σ values and averaged across 100 repetitions. **C, F** Differences in mean accuracies between the models (T-TVFC – Covariance-based HMM).

In the second scenario, *X* was directly obtained from 11 network timeseries obtained from resting-state 7T fMRI data. **Figure 2E** shows the mean classification accuracy scores of each model across 100 repetitions for each combination of *p* (where *p* timeseries were randomly sampled) and σ (from 0.001 to 0.18). Again, T-TVFC consistently outperformed the covariance-based HMM in capturing the interactions of interest, as evidenced in **Figure 2F**. As in the first scenario, the covariance-based HMM can sometimes achieve comparable or even higher accuracy to the T-TVFC model under conditions of very low noise levels and only one or two predictors. However, this advantage diminishes as noise levels increase.

Overall, these simulations show that T-TVFC, being more specifically tailored to capture the interaction between *X* and *Y*, is more effective at characterising the brain-behaviour association under study when our hypothesis about the relevant connections involved is correct.

Additionally, we evaluated the performance of T-TVFC against a clustering-based approach applied to SWC matrices. Simulated X and Y time series (with σ = 0.1) were concatenated channel-wise into a single matrix Z. A k-means clustering algorithm with K=7 states was applied to the windowed-FC matrices. This was performed in two ways: one using all *p*×*p* region pairs, and the other focusing on the targeted *p*×*q* pairs. We evaluated it across different values of *p* and varying window sizes. The state time courses inferred by each model were used to predict trial conditions over time via logistic regression (baseline accuracy = 0.5). None of the sliding-window configurations outperformed T-TVFC in predicting trial conditions. These findings are presented in **Figure S5**.

### 3.2. REAL DATA

Next, we applied the T-TVFC model to 7T task fMRI data from 60 subjects to investigate the thalamocortical connections associated with distinct cognitive processes involved in reasoning. We compared the performance of this model with that of the covariance-based HMM. Both models were configured with *K* = 3 states, as we observed that more states did not offer additional task-related information (refer to **Figure S6** for the results with *K* = 4 states). In the T-TVFC model, each state represents a distinct pattern of thalamocortical FC, modelled using a regression approach where 11 cortical timeseries are used as regressors (*X*) and 3 thalamic timeseries as the response (*Y*). As before, in the covariance-based HMM, both sets of timeseries were treated collectively as *Z*, and characterized by state-dependent covariances modelling connections across all pairs of regions.

We assessed each model’s performance using ordinal logistic regression, given the ordered nature of the conditions (binary, ternary and quaternary). A perfect classification corresponds to an accuracy of 1, while 1/3 indicates random guessing among three conditions.

As depicted in **Figure 3A**, the T-TVFC model achieved higher accuracy in discriminating levels of complexity compared to the covariance-based HMM. This suggests that focusing on specific connections enhances the model’s ability to capture the neural dynamics associated with varying levels of problem-solving difficulty. Notably, the T-TVFC model appears to detect early complexity-based dynamics that remain hidden in the conventional HMM, possibly because, at the beginning of a trial, it is mostly the corticothalamic connections the ones that exhibit a task effect. In contrast, the conventional HMM, which considers connectivity across and within all thalamic and cortical regions, picks up changes later in the trial when a broader range of connections becomes involved in supporting the behaviour, leading to poorer early-in-the-trial accuracy in classifying cognitive demand.

**Figure 3.**
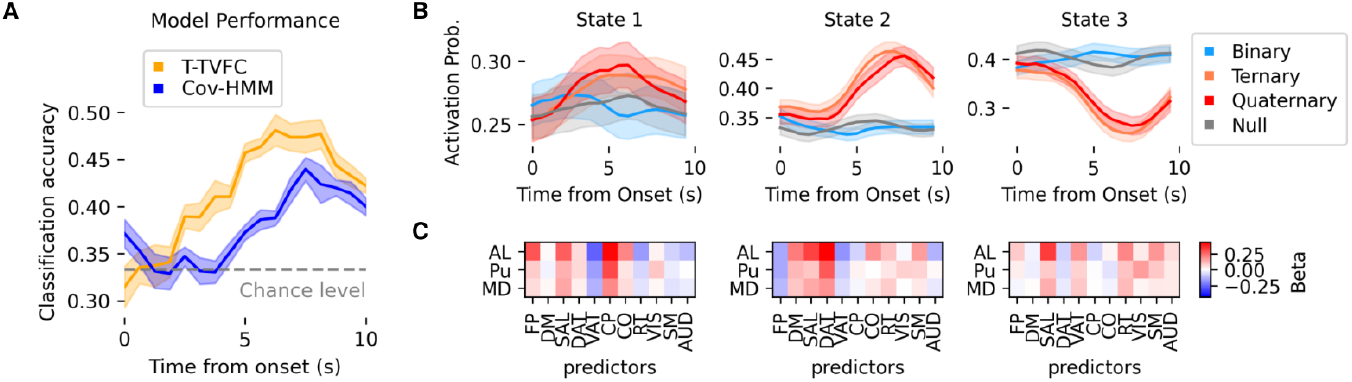
Model performance in classifying task conditions using real data. **A** Performance of the T-TVFC (orange) compared to the covariance-based HMM (blue) over time. Accuracy values (mean and standard deviation across 10 repetitions) are obtained from ordinal logistic regression and evaluated across complexity levels (binary, ternary, and quaternary), compared against the 1/3 chance level. **B** Trial-locked (mean and standard error of the mean) state time courses from T-TVFC for null (grey), binary (blue), ternary (orange) and quaternary (red) conditions. **C** Each state corresponds to a distinct set of β coefficients. The y-axis represents three thalamic nuclei, while the x-axis represents the 11 functional networks acting as predictors.

We further compared the performance of T-TVFC with the SWC approach, applied to both all *p*×*p* region pairs and targeted *p*×*q* pairs. Rectangular windows ranging from 4 to 10 TRs were used, with a one-TR step size between shifts. Across all configurations, the SWC approach consistently underperformed compared to T-TVFC in predicting trial conditions (**Figure S7**).

In addition to evaluating the overall performance of the T-TVFC model, we looked at the state-specific information related to the task, aiming to interrogate the spatiotemporal characteristics of thalamocortical connectivity patterns associated with varying cognitive demands. We can visually appreciate the distinct activation probabilities, particularly when comparing ternary and quaternary trials to binary trials. As observed, binary trials resembled null trials, where participants simply pressed a designated button without engaging in cognitive processing. This similarity is not surprising given that participants’ performance in binary trials was nearly identical to their performance in null trials (see description of the data in *Methods*). The trial-locked trajectories of each state across conditions are shown in **Figure 3B**. There is a clear pattern of increased activation in state 2 and deactivation in state 3 during complex trials (ternary and quaternary). These effects were particularly prominent approximately five seconds after the presentation of the trial. The mirrored trajectories of the two states suggest a dynamic interplay between the thalamus and key brain networks. Specifically, during complex trials, there is a direct relationship between the thalamus and the DM and DAT networks, coupled with an inverse relationship between the thalamus and the FP and VAT networks. As the trials progressed, these trajectories gradually returned to baseline, indicating a reconfiguration of thalamocortical interactions during later stages of problem-solving, possibly reflecting adaptive neural processes. The regression coefficients for each state are illustrated in **Figure 3C**.

## 4. DISCUSSION

The present study introduces T-TVFC, a novel approach to explore the temporal changes of specific brain connections, driven by previous hypotheses. Unlike covariance-based models, which are based on correlations across all variable pairs and risk diluting the contributions of individual connections, T-TVFC isolates specific subsets of connections. It achieves this by performing separate regressions for each variable in one set (acting as the response) against the variables in another set (acting as predictors). This targeted approach enhances the ability to examine how particular connections behave in relation to the cognitive construct in question. It is important to recognize that connectivity estimates, including those derived from T-TVFC, remain partially influenced by signal amplitude. Higher amplitudes improve the signal-to-noise ratio and may bias the connectivity estimates—a challenge shared by most connectivity-based approaches. Nevertheless, T-TVFC outperforms methods such as the HMM covariance approach and sliding-window techniques, enabling the detection of nuanced connectivity dynamics that broader covariance-based or less temporally resolved methods might overlook. Importantly, even though the model allows for analyses that are more hypothesis-driven than some existing alternatives, it should be kept in mind that it is a data analysis technique and therefore stays at a phenomenological level (Vidaurre, 2024).

To showcase the method, we applied it to investigate thalamocortical interactions during a reasoning task. Here, T-TVFC captured fluctuations in thalamocortical connectivity that hold a stronger association with task-related cognitive demands than an HMM that considers connectivity across all pairs of regions. This finding corroborates previous research highlighting the thalamus’s pivotal role in coordinating the task-induced activity of large-scale brain networks (Shine, 2021). By isolating thalamocortical connectivity, T-TVFC allowed us to attribute observed task-related dynamics specifically to these connections, avoiding the ambiguity introduced by whole-brain covariance analyses, where numerous connections— including task-irrelevant ones—might drive observed effects. While recent advances, such as the whole-brain dynamic FC atlas (Peng et al., 2023), provide a comprehensive map of brain-wide interactions, T-TVFC offers a more focused, hypothesis-driven framework for investigating specific connectivity patterns. By concentrating on targeted thalamocortical dynamics, T-TVFC complements these broader approaches, providing a detailed understanding of the task-related interactions that might be obscured in large-scale analyses.

Moreover, T-TVFC’s focus on specific connections provides an advantage in scenarios where different subsets of connections may vary independently over time. In traditional state-based approaches, like the HMM, a single sequence of states is used to describe the data, meaning that the spatial and temporal properties of those states apply to all connections (either *p*×*p* or *p*×*q*, in this case). That is, if two sets of connections fluctuated independently, these models would require additional states to capture all possible configurations. For example, if we were modelling two connections that alternate between positive and negative coupling, and the timing of the switches between positive and negative is independent for the two connections, we would need four distinct states to represent all combinations. This limitation applies to the HMM, T-TVFC, and sliding-window approaches. However, T-TVFC can ameliorate this issue by allowing separate analyses for each subset of connections.

Another potential advantage of T-TVFC lies in its capacity to model directionality, compared to the symmetric focus of the covariance-based approach, which uses bidirectional correlations or covariances across all pairs of variables. In contrast, T-TVFC, by design, emphasizes the directional influence of one set of variables (predictors) on another set (response), thereby capturing specific and potentially non-reciprocal interactions. This directionality may allow for a nuanced exploration of how specific variables drive changes in others, offering insights that may be obscured when exclusively examining symmetrical correlations. In principle, this approach could be used to quantify information flow. However, fMRI poses specific limitations in this regard because it is an indirect measurement of neural activity via blood flow changes and the regions’ vascular dynamics might be different, therefore making a strict interpretation of directionality cumbersome. Future applications of T-TVFC to direct neural recordings such as magnetoencephalography (MEG) could provide directional insights into neural interactions.

In conclusion, this study demonstrates the effectiveness of T-TVFC, for instance in revealing the unique temporal dynamics supporting the execution of a complex reasoning task. By capturing specific neural interactions, T-TVFC can be a useful tool for disentangling the various concurrent threads of processing in the brain.

## AUTHOR CONTRIBUTIONS

SA and DV: Conceptualization, Methodology, Formal Analysis, Writing – original draft. LJH: Data collection and preprocessing. SA, DV, LJH, JMS, LC: Writing – review & editing.

## DATA AND CODE AVAILABILITY

Data is available at https://github.com/macshine. Code is openly available at https://github.com/sonsolesalonsomartinez/TTVFC

## Supplementary Material

**Figure S1.**
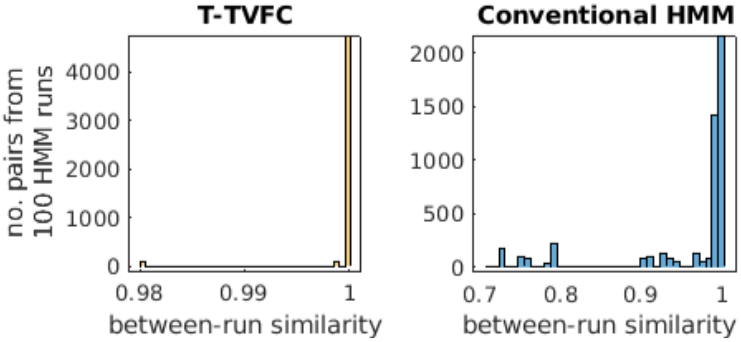
Similarity scores across 100 repetitions of the T-TVFC and the conventional (covariance-based) HMM.

**Figure S2.**
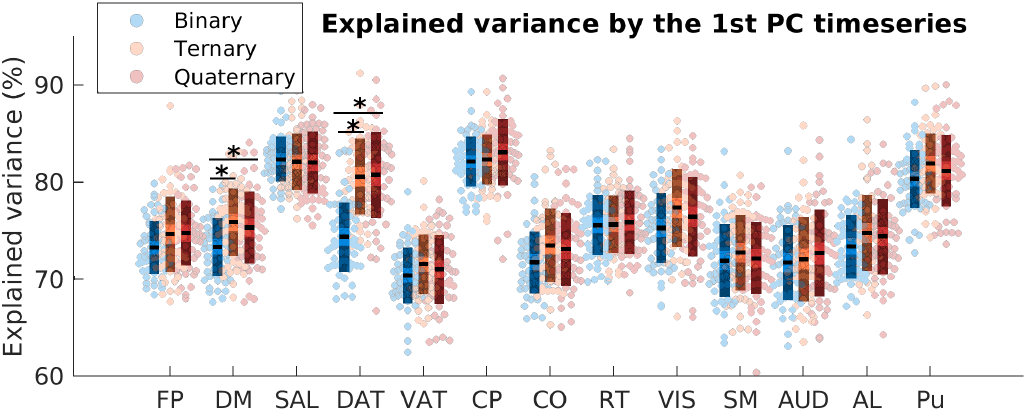
Percentage of explained variance by the,rst principal component for the 11 cortical networks and two thalamic nuclei (AL and Pu). PCA was not applied to MD nuclei as it consists of only one region. Individual scores represent trial-averaged for each subject and per condition (binary, ternary and quaternary). Boxes represent the standard deviation, lighter shading indicates the standard error, and the dashed line denotes the mean across subjects. * indicates pairwise differences with FWE-corrected p-values < 0.05, obtained through non-parametric (n=5000) permutation testing. Abbreviations: AL: Anterolateral thalamic nuclei; AUD: Auditory; CP: Cinguloparietal; CO: Cingulopercular; DM: Default Mode; DAT: Dorsal Attention; FP: Frontoparietal; MD: Medial Dorsal thalamic nuclei; Pu: Pulvinar thalamic nuclei; RT: Retrosplenial Temporal; SAL: Salience; SM: Sensorimotor; VAT: Ventral Attention; VIS: Visual.

**Figure S3.**
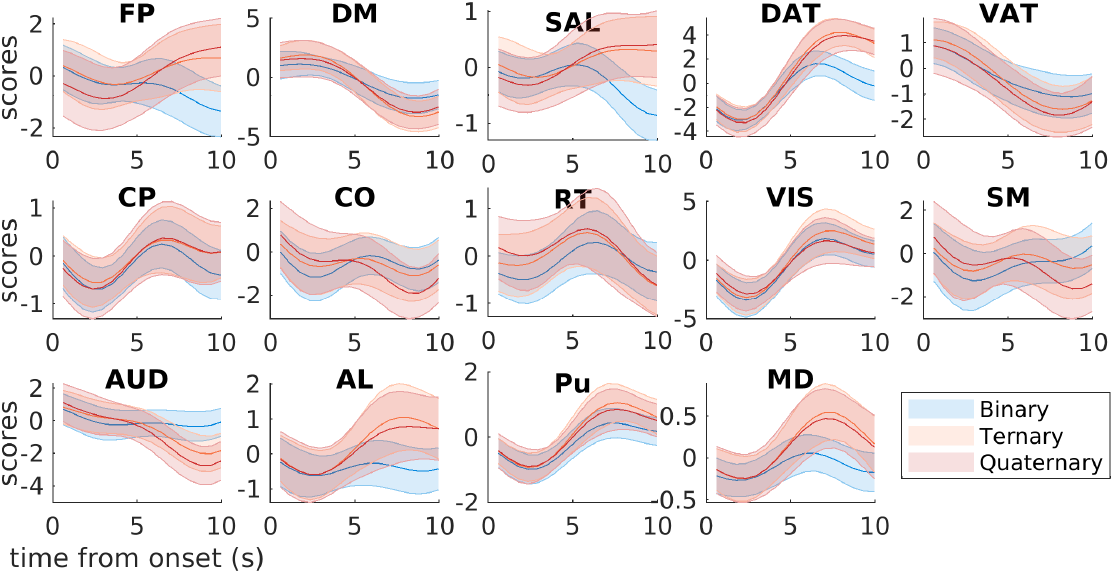
Mean (± SD) trial-locked time series across subjects from 11 cortical networks and three thalamic nuclei under binary, ternary and quaternary conditions. Abbreviations: AL: Anterolateral thalamic nuclei; AUD: Auditory; CP: Cinguloparietal; CO: Cingulopercular; DM: Default Mode; DAT: Dorsal Attention; FP: Frontoparietal; MD: Medial Dorsal thalamic nuclei; Pu: Pulvinar thalamic nuclei; RT: Retrosplenial Temporal; SAL: Salience; SM: Sensorimotor; VAT: Ventral Attention; VIS: Visual.

**Figure S4.**
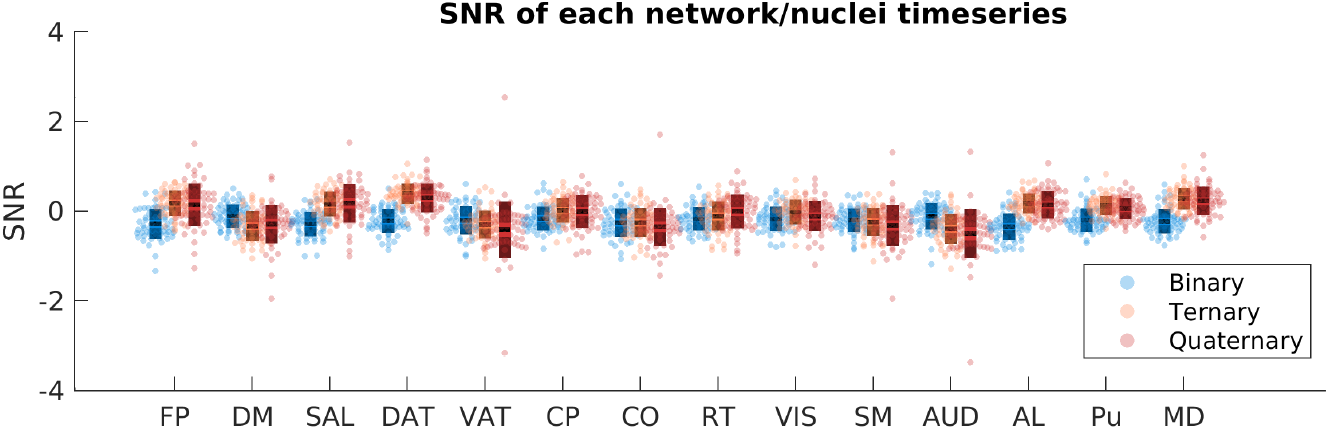
Signal-to-noise ratio (SNR) for the 11 cortical network timeseries and three thalamic nuclei timeseries. Individual scores represent trial-averaged for each subject and per condition (binary, ternary and quaternary). Boxes represent the standard deviation, lighter shading indicates the standard error, and the dashed line denotes the mean across subjects. Abbreviations: AL: Anterolateral thalamic nuclei; AUD: Auditory; CP: Cinguloparietal; CO: Cingulopercular; DM: Default Mode; DAT: Dorsal Attention; FP: Frontoparietal; MD: Medial Dorsal thalamic nuclei; Pu: Pulvinar thalamic nuclei; RT: Retrosplenial Temporal; SAL: Salience; SM: Sensorimotor; VAT: Ventral Attention; VIS: Visual.

**Figure S5.**
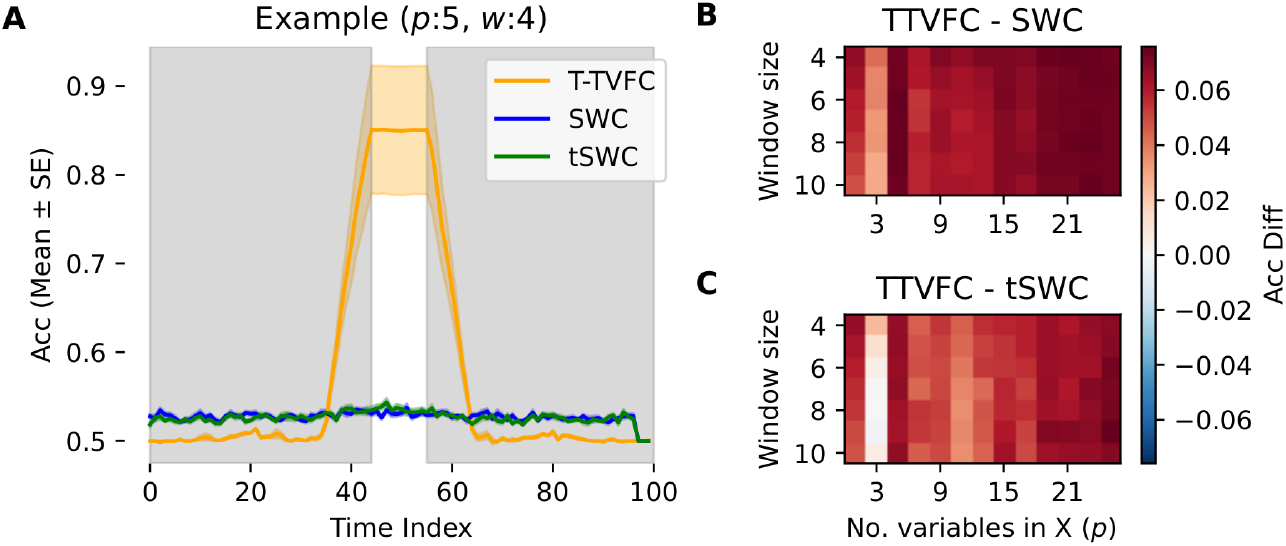
Model performance compared to SWC approaches using simulated data. **A**. Example illustrating the performance of the T-TVFC (orange) compared to all-pairs SWC (in blue) and speci,c-pairs SWC (tSWC, in green) over time for a speci,c parameter con,guration (*p* = 5, σ = 0.1 and *w* = 4). Accuracy in predicting trial conditions, represented as the mean ± standard error over 10 repetitions, was calculated using logistic regression and evaluated against the chance level of 0.5. The white patch indicates the period where the simulated task-dependent state was active. **B**. Differences in mean accuracies between T-TVFC and all-pairs SWC. **C**. Differences in mean accuracies between T-TVFC and speci,c-pairs SWC, across varying values of *p* and *w*. Mean accuracies for each model were computed during the interval when there are differences between conditions. *p*: number of predictors in X. σ: levels of noise. *w*: window size used in SWC.

**Figure S6.**
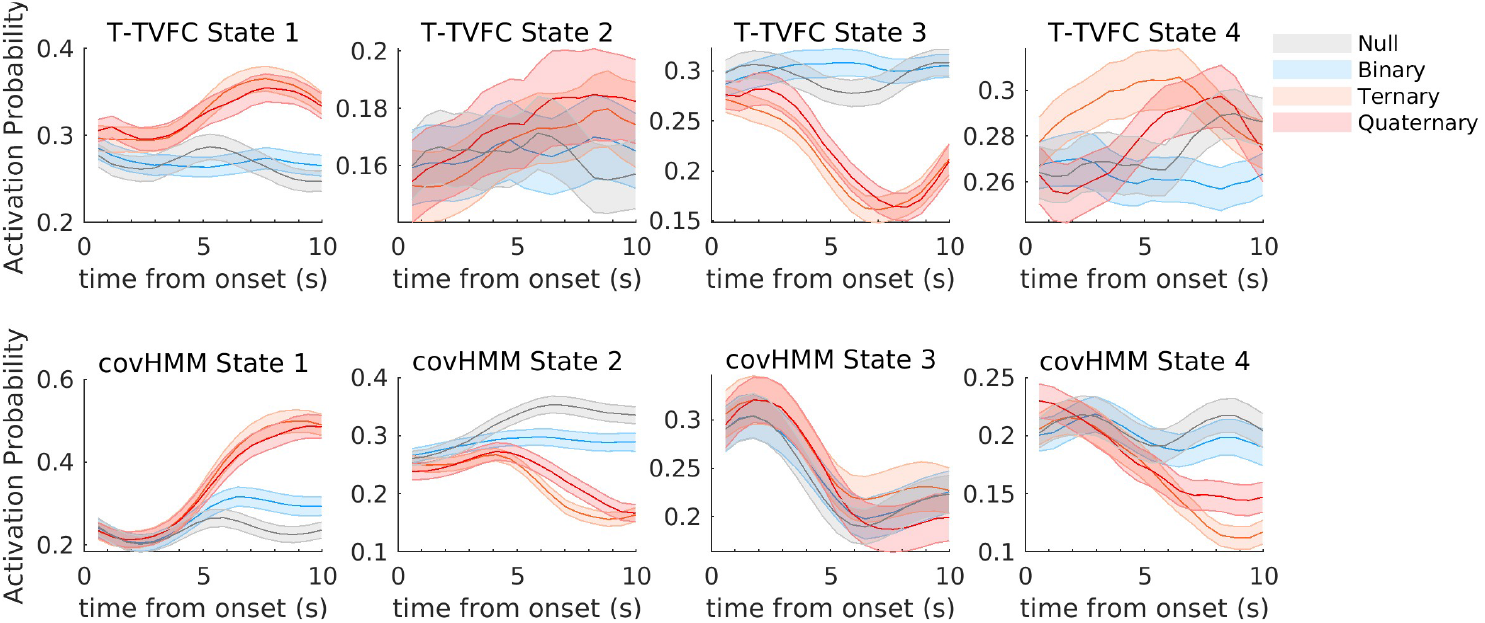
Trial-locked state time courses from T-TVFC (*top*) and covariance-based HMM (*bottom*), both configured with *K* = 4 states. Increasing the number of states from *K* = 3 to *K* = 4 does not provide additional insights but instead divides existing states into more granular representations.

**Figure S7.**
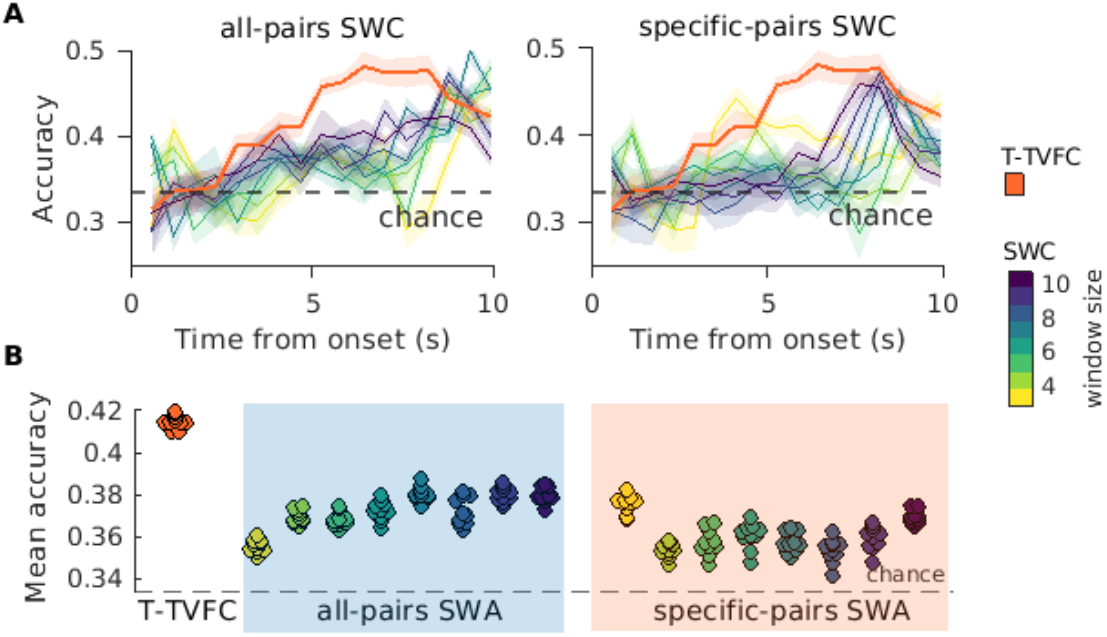
Model performance compared to SWC approaches using real data. **A**. Classi,cation accuracy of T-TVFC (orange) compared to all-pairs SWC (leD) and speci,c-pairs SWC (right) across a range of window sizes (3 to 10 TRs; see color bar). Accuracy values, represented as the mean and standard deviation over 10 repetitions, were obtained using ordinal logistic regression and evaluated at different levels of task complexity (binary, ternary, and quaternary classi,cation). Results are compared against the 1/3 chance level. **B**. Average classi,cation accuracy over the 10-second trial duration for T-TVFC (orange) compared to all-pairs SWC (blue shading) and speci,c-pairs SWC (orange shading), evaluated across varying window sizes.

